# Novel regimens for treatment of *Mycobacterium avium* lung disease based on advanced *in vitro* systems and the mathematics of basis functions

**DOI:** 10.64898/2026.03.30.715241

**Authors:** Shashikant Srivastava, Sanjay Singh, Gunavanthi D Boorgula, Pamela J McShane, Tawanda Gumbo

## Abstract

Azithromycin plus ethambutol plus rifabutin (azithromycin-ethambutol-rifabutin) is the standard-of-care (SOC) for *Mycobacterium avium*-complex lung disease. The SOC achieves sustained sputum culture conversion in only 43-53% of patients, after an average of 18 months of therapy. Recent quantitative analyses ranked omadacycline, ceftriaxone, and minocycline highest for microbial kill. Azithromycin-minocycline-ethambutol, azithromycin-omadacycline-ethambutol, epetraborole-omadacycline-ethambutol, ceftriaxone-omadacycline-rifabutin, and the SOC were compared in the intracellular hollow fiber system model of *M. avium* lung disease (HFS-MAC). HFS-MAC units were treated once daily for 28 days to mimic the intrapulmonary pharmacokinetics of each drug. The ceftriaxone concentrations measured in the HFS-MAC were only 1% of those achieved in the lung by standard clinical doses. Changes in the bacterial burden were described using basis functions (BF). For liquid cultures, BF 1 (BF_1_) was described by a linear regression-based slope, with steepest kill slope (95% Confidence interval) of 7.87 (1.52 to14.23) by ceftriaxone-omadacycline-rifabutin versus 1.04 (-0.84 to 2.92) for SOC. For the CFU/mL readout, the BF_1_ steepest non-linear kill slope was for ceftriaxone-omadacycline-rifabutin of 0.55 (0.35 to 0.98) log_10_ CFU/mL/day versus 0.16 (0.07 to 0.25) log_10_ CFU/mL/day for the SOC. Thus, ceftriaxone-omadacycline-rifabutin is potentially better than the SOC, even though further ceftriaxone dose optimization is required. BF_2_ described rebound growth and drug-resistant subpopulation growth, and demonstrated that contrary to popular belief, SOC rebound was best explained by ethambutol-resistance (r^2^>0.99, *p*=0.01) and not by azithromycin-resistance (r^2^=0.27, p=0.32), questioning ethambutol’s role in the SOC. The BF framework is potentially easy to adapt for modeling other anti-infective agents across many infectious diseases.

## INTRODUCTION

*Mycobacterium avium complex* (MAC) lung disease is treated with the standard-of-care (SOC) that is a combination of azithromycin (or clarithromycin) plus rifabutin (or rifampin) plus ethambutol (1). The macrolide performs the microbial kill (i.e., main drug), while ethambutol and rifabutin prevent emergence of resistance to the macrolide (2–5). Thus, macrolide-resistance drives failure of the combination therapy. Consequently, the only susceptibility testing believed to matter in the clinic is that of the macrolides. Meta-analyses have demonstrated that SOC was associated with a 6-month sustained sputum culture conversion (SSCC) in 43-53% of patients, despite an average therapy duration 18 months (1, 6, 7). This response rate with a prolonged therapy duration is undoubtedly suboptimal. Moreover, in a review of 364 patients with MAC lung disease treated the SOC, 20% developed hepatotoxicity, 20% leukocytopenia, 29% thrombocytopenia, and 9% cutaneous reactions, while 8% developed ocular toxicity (8). Thus, newer combination therapies with less toxicity, better efficacy, and shorter duration are the need of the hour.

A recent quantitative analysis of pre-clinical pharmacokinetics (PK)-pharmacodynamics (PD) models demonstrated that 87% of the studies were performed using the intracellular hollow fiber system model of MAC lung disease (HFS-MAC), and 13% of studies in the mouse model (9). The study also introduced a quantitative quality score for experiments. Second, it introduced quantitative methods to compare the effect of single drugs, as well as of combination therapy, based on how well drugs kill MAC below day 0 (pre-treatment) bacterial burden (*B_0_*) at optimal dose (9). *B_0_* is one of the best predictors of therapeutic outcomes in patients, consistent with the inoculum effect (6, 7, 10, 11). Kill below *B_0_* was then standardized to the SOC kill below *B_0_*, to give an index of fold-kill compared to the SOC. The highest ranked drugs were omadacycline (69-fold), tedizolid (19-fold), ceftriaxone (8-fold), and minocycline (>3-fold) (9). Here, we tested several combinations of these highest ranked drugs, to identify the best combination to advance for further testing.

When using the CFU/mL readout, microbial kill followed by rebound due to resistance emergence creates a complex function. In order to compare the composites of speeds of kill and extent of kill as well for rebound growth due to resistance-emergence, and to integrate the different phases of microbial effects of different combinations – kill, resistance emergence, and rebound growth, we introduce the concept of “basis functions” (BF) (12, 13). BF 1 (BF_1_) describes the kill curve, and BF_2_ the rebound and resistance emergence, and the composite function (*f*_C_) is defined as follows:

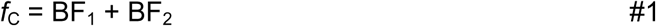

The kill curve for BF_1_ has been described elsewhere, based on the ordinary differential equation (ODE) that describe the dynamics of MAC drug susceptible (wild type, *w*) and isogenic resistant strains (*m*), which is the difference between a growing bacterial population (growth rate – described by a logistic function) and microbial kill, as follows (14–20):

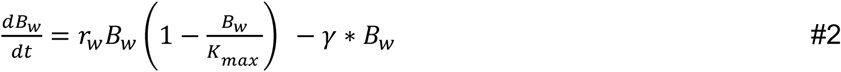

𝜸-slope starts at time of 0 (x_t,0_) and ends at the time 1 (x_t,1_) when rebound (therapy failure due to resistance emergence) starts, termed hinges. In other words, BF_2_ is when there is no microbial kill, that is γ = 0, so that based on the ODE the expression is now the standard logistic growth function, 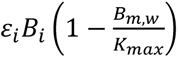 with growth rate represented by 𝜀_i_. BF_2_ 𝜀 starts at the hinge x_t,1_and ends at the end of the experiment at time 2 (x_t,2_). The hinge is defined by (x, y) coordinates, that is time in days, and bacterial burden. As regards to time-to-positivity in Mycobacterial Growth Indicator Tube (MGIT) liquid cultures, the BFs are described by linear regression slopes, that cross at the hinge co-ordinates. The BFs are thus different from those for CFU/mL but can be integrated into the same equation #1. Here we demonstrate how this approach can be used to identify the best combination regimens.

## RESULTS

### Quality control of the HFS-MAC study

The quality control (QC) score for the HFS-MAC study was 14 out of total possible of 20 (9). This places it in the “adequate” category. The percentage coefficient of variation (%CV) between replicates (average of all sampling days) for bacterial burden for log_10_ CFU/mL and TTP are shown in **Figure 1A** and **Figure 1B**. These show that the average %CV for each pair of HFS-MAC for a given drug combination was below 25%. **Figure 1C** shows %CV data for drug concentration measurements, for each drug. The median score for each drug was below 25%. **Figure 1D** shows the summary for all replicate HFS-MAC units, expressed as medians and 95% Confidence Intervals (CI). These results mean the experiments passed the QC based on both % CV and quality score **(Table 1)**.

**Figure 1.**
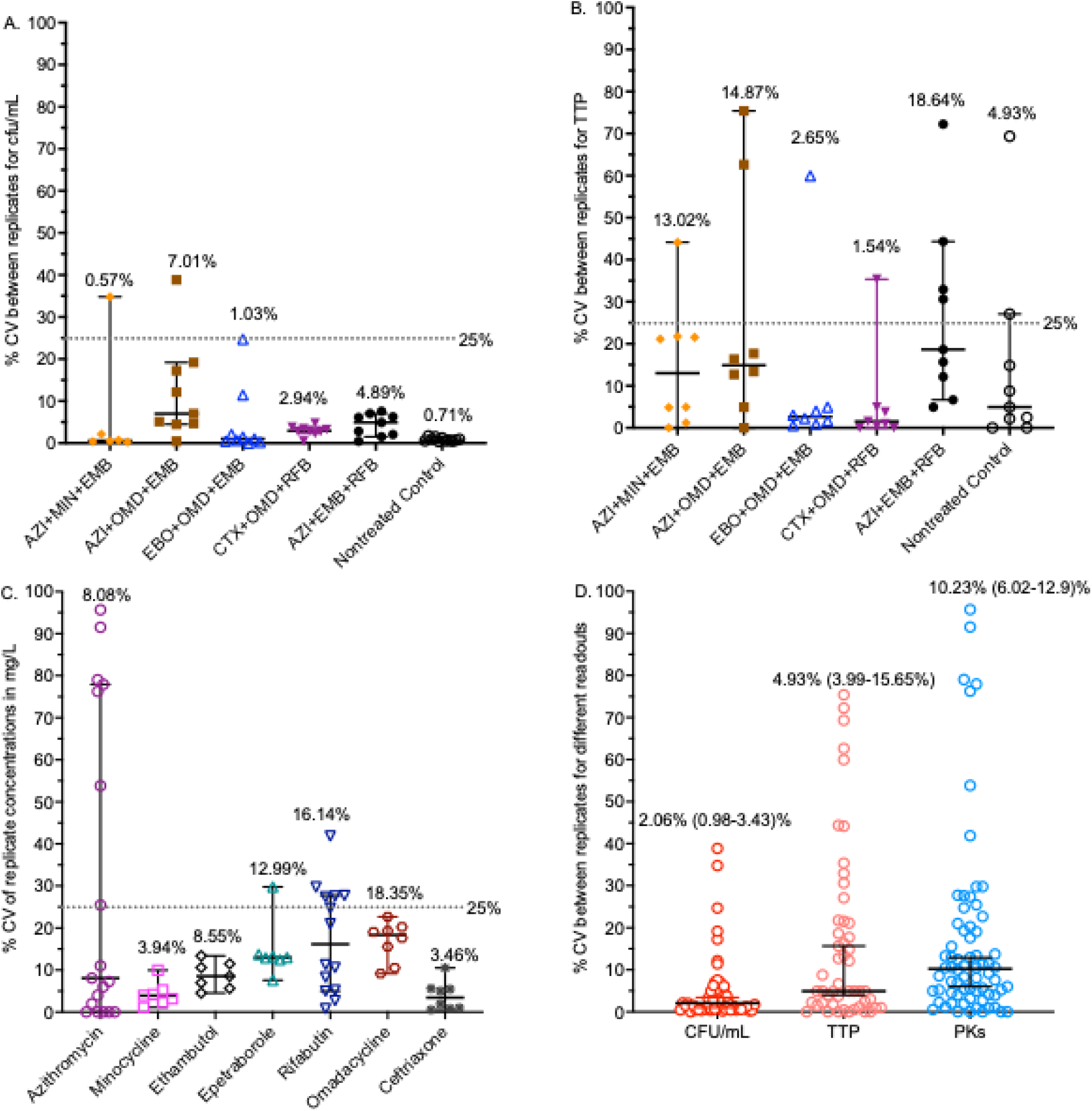
Coefficient of variation between HFS-MAC replicates for use as quality control. Bars are medians, and error bars are 95% confidence intervals. Data points are for each pair of hollow fiber system units at each sampling point. AZI=azithromycin, CTX=ceftriaxone, EMB=ethambutol, EBO=epetraborole, MIN=minocycline, OMD=omadacycline, RFB=rifabutin. **A.** Percent coefficient of variation (CV) between replicates for the log_10_ CFU/mL replicates. **B.** CV for time-to-positivity (TTP) readout between replicates. **C.** CV between replicates for drug concentrations. **D.** All readouts combined, numbers are point estimates and brackets are 95% confidence intervals.

**Table 1.** Quantitative Quality Criteria Performance and Score in the combination HFS-MAC study.

| Criteria | Passed |
| --- | --- |
| Standard of care arm | Yes, tested |
| Standard of care arm, $\gamma$ 95% interval versus historical controls | Yes, compared to historical controls in prior publication (16) |
| 5 conditions/dosing | No |
| 5 MAC isolates | No |
| 5 Monte Carlo experiment doses | Yes |
| 25% Coefficient of variation between replicates | Yes, for PKs and bacterial burden readouts |
| Quality score | Yes, Adequate |

### PK findings and concentrations achieved in HS-MAC

The measured drug concentration-time profiles in HFS-MAC units and PK model derived concentration-time profiles are shown in **Figure 2**. The PK model diagnostics are shown in **supplementary Figures S1 and S2**. This means that there was good model fit, minimal bias, and parameter estimation was accurate. PK parameter estimates from the modeling and the PK/PD exposure estimates (incorporating the MICs) were as shown in **Table 2**.

**Figure 2.**
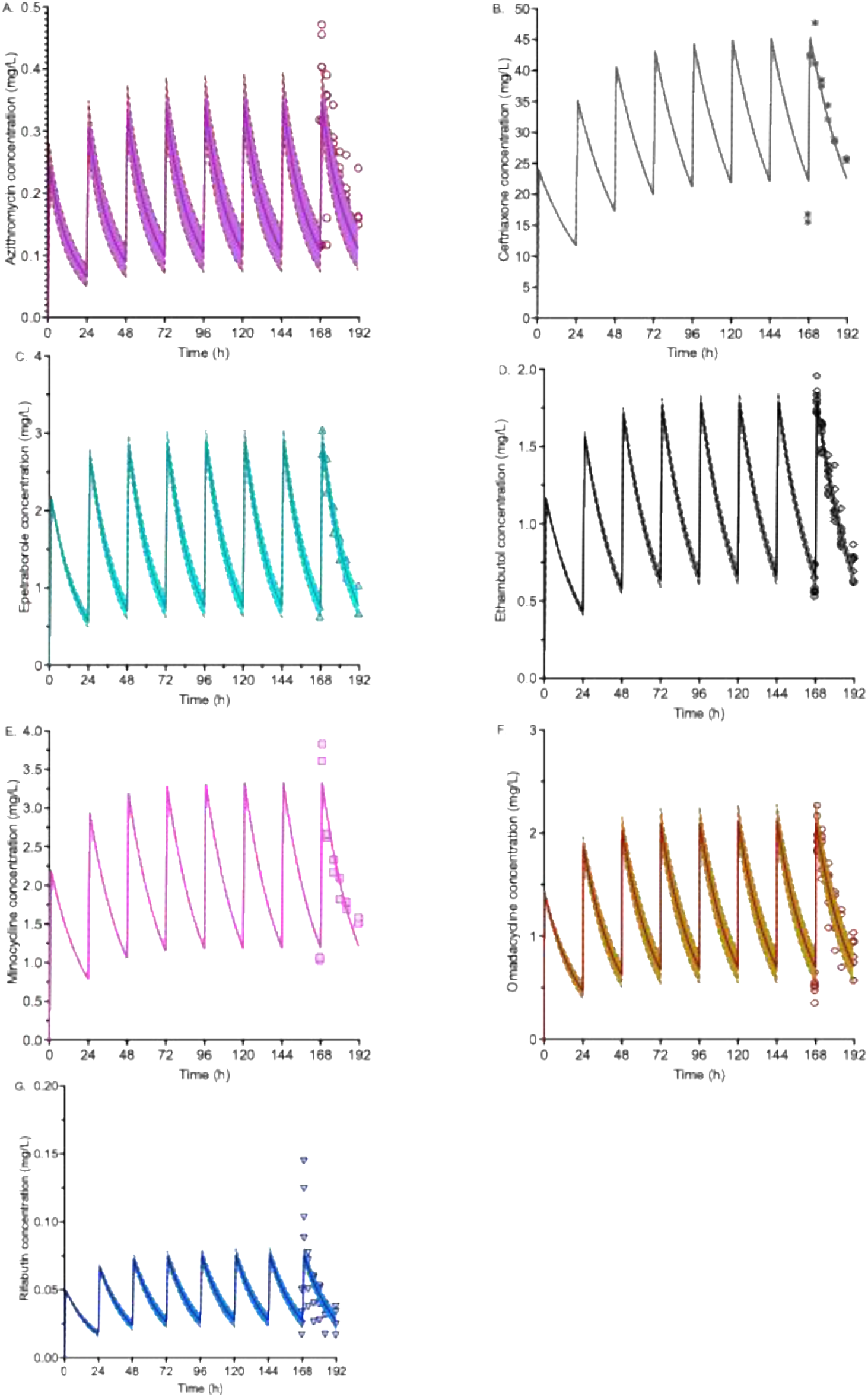
Drug concentration-time profiles based on measurements in the HFS-MAC. Shading is for 95% confidence intervals of PK modeled concentrations; in several of these the confidence intervals were very narrow as to appear like a single line. Symbols are the measured drug concentration replicates from samples on day 7 (168h). **A**. Azithromycin. **B**. Ceftriaxone. **C.** Epetraborole. **D.** Ethambutol. **E.** Minocycline. **F.** Omadacycline. **G.** Rifabutin.

**Table 2.** Pharmacokinetic parameter estimates for drug concentration-time profiles measured in the HFS-MAC.

| Drug | Clearance (L/h) |  | Volume (L) |  | AUC<sub>0-24</sub> (mg*h/L) |  | AUC<sub>0-24</sub>/MIC | Peak concentration (mg/L) |  |
| --- | --- | --- | --- | --- | --- | --- | --- | --- | --- |
|  | Estimate | SD | Estimate | SD | Estimate | SD |  | Estimate | SD |
| **Azithromycin** | $5.68 \times 10^{-2}$ | $3.63 \times 10^{-3}$ | 0.97 | 0.40 | 4.93 | 0.05 | 0.62 | 0.35 | 0.11 |
| **Ceftriaxone** | $1.15 \times 10^{-2}$ | $4.81 \times 10^{-5}$ | 0.37 | $6.12 \times 10^{-3}$ | 691.25 | 0.01 | 172.82 | 44.65 | 0.85 |
| **Epetraborole** | $3.2 \times 10^{-2}$ | $3.32 \times 10^{-3}$ | 0.54 | $3.68 \times 10^{-3}$ | 36.78 | 0.11 | 36.78 | 2.90 | 0.16 |
| **Ethambutol** | $1.85 \times 10^{-2}$ | $1.06 \times 10^{-3}$ | 0.42 | $3.14 \times 10^{-3}$ | 25.38 | 0.03 | 3.17 | 1.79 | 0.06 |
| **Minocycline** | $2.5 \times 10^{-2}$ | $1.15 \times 10^{-4}$ | 0.56 | $7.49 \times 10^{-3}$ | 46.78 | $2.51 \times 10^{-4}$ | 46.78 | 3.33 | 0.01 |
| **Omadacycline** | $2.13 \times 10^{-2}$ | $2.67 \times 10^{-3}$ | 0.46 | $1.69 \times 10^{-2}$ | 28.65 | $7.83 \times 10^{-2}$ | 7.16 | 2.12 | 0.14 |
| **Rifabutin** | $4.81 \times 10^{-1}$ | $9.75 \times 10^{-2}$ | 10.6 | $2.3 \times 10^{-1}$ | 1.06 | $4.81 \times 10^{-3}$ | 8.48 | 0.08 | $8.32 \times 10^{-3}$ |

### Microbial kill modeling using MGIT liquid culture readout

Liquid culture derived time-to-positivity (TTP)-based microbial kill curves during 42 days of study are shown in **Figure 3A**. The higher the TTP, the lower the bacterial burden. No growth was modeled as 57-days since cultures were declared negative after 56 days of time-in-protocol (incubation). The first comparison was of maximal microbial kill calculated as highest TTP minus *B_0_* (or amplitude) by regimens on day 7. **Figure 3A** shows that the highest amplitude was for ceftriaxone-omadacycline-rifabutin and epetraborole-omadacycline-ethambutol combinations. To put this into context, we calculated how much better the novel regimens amplitude was compared to the SOC’s amplitude, for the TTP readout. The mean (± standard deviation) fold-SOC was 2.82±2.64-fold for azithromycin-minocycline-ethambutol, 4.21±0.66-fold for azithromycin-omadacycline-ethambutol, and 4.68±0.01-fold for either the ceftriaxone-omadacycline-rifabutin or the epetraborole-omadacycline-ethambutol regimens.

**Figure 3.**
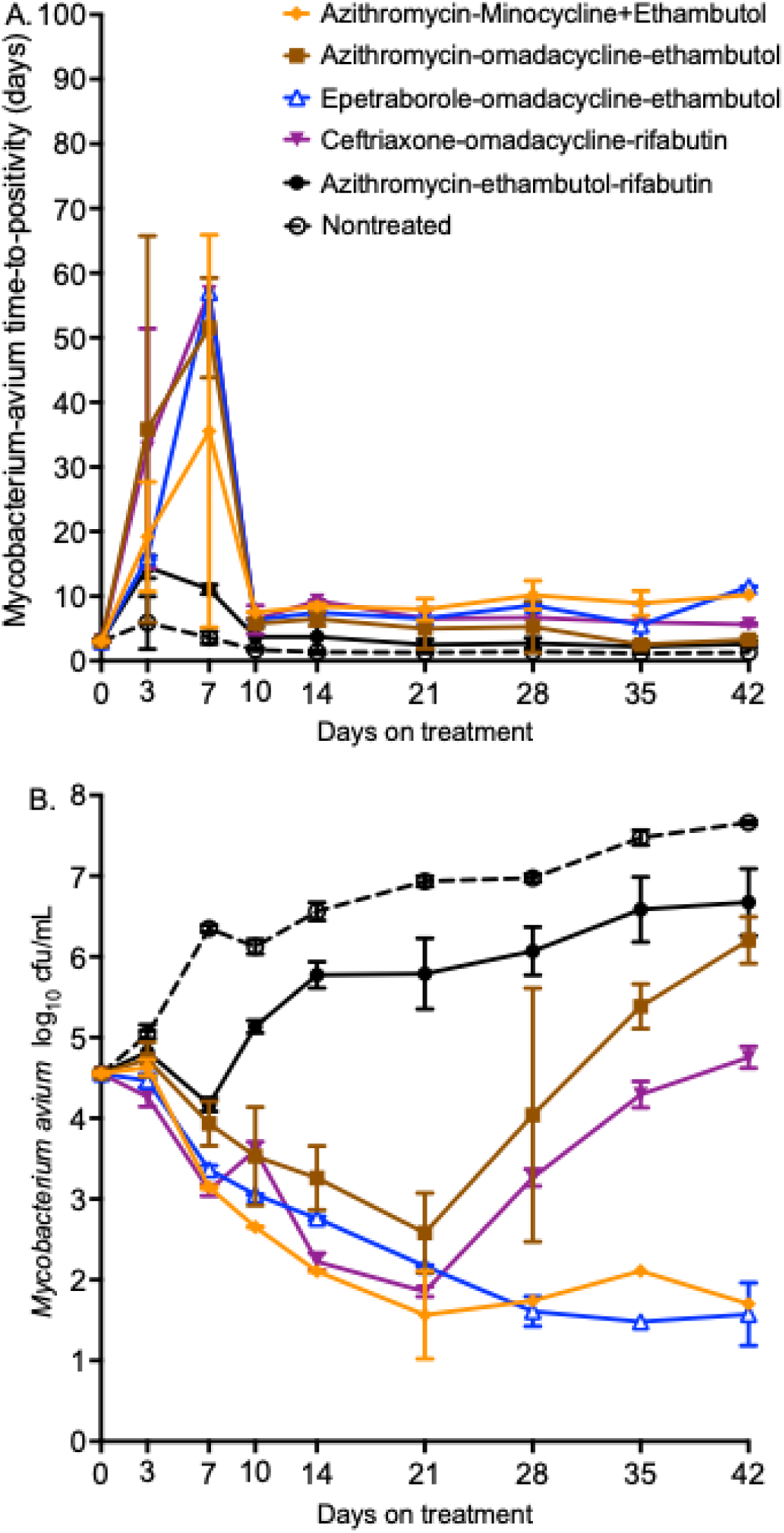
Pharmacodynamics of the different combination in HFS-MAC using TTP and CFU/mL readouts. Symbols are mean, and error bars are standard deviations. **A.** Time-to-positivity in the Mycobacteria Growth Indicator Tube assay. **B.** Colony forming units on agar

**Figure 3A** also shows that all regimens had a triphasic pattern of effect, which means the regimens could be described using three linear BFs. Non-treated controls were described by only one BF. **Table 3** shows the three BFs and slopes. Since the bacterial burden TTP units were in units of days and X-axis is also in days, the slopes are unitless. For BF_1_ the slope for the SOC was 1.04 with a bacterial burden (TTP) of 11.06 at hinge 1, while the non-treated controls were -0.05 with hinge at 42 days and TTP of 1.29. The steepest BF_1_ slope was for ceftriaxone-omadacycline-rifabutin, which was 7.57 times faster than the SOC, with 95%CIs that did not overlap with any other regimen except azithromycin-minocycline-ethambutol. Hinge 1 on day 7 shows that the lowest bacterial burden (highest TTP) at the end of kill phase was with ceftriaxone-omadacycline-rifabutin at 57 days. For BF_2_, the 95%CIs of the slopes crossed zero for azithromycin-minocycline-ethambutol and ceftriaxone-omadacycline-rifabutin, which means the rebound slope was statistically zero for these two regimens. BF_3_ demonstrated a zero slope for all regimens because the 95%CI of all crossed zero, which means there was no change in bacterial burden as time changed for each treatment regimen.

**Table 3.** Changes in bacterial burden for different treatment regimens based on liquid cultures florescence readout.

| Regimen | $B_0$: days | Hinge 1 | Slope 1 for BF1 | Hinge 2 | Slope 2 for BF2 | Hinge 3 | Slope 3 for BF3 |
| --- | --- | --- | --- | --- | --- | --- | --- |
|  |  | day, day | [unitless] | [day, day] | [unitless] | [day, day] | [unitless] |
| Azithromycin-minocycline-ethambutol | 3.02 [0 - 21.27] | 7, 35.54 | 4.62 [-1.63 - 10.01] | 10, 7.53 | -9.34 [-40.12 – 21.44] | 42, 10.2 | 0.06 [-0.07 - 0.19] |
| Azithromycin-omadacycline-ethambutol | 3.02 [1.68 - 4.35] | 7, 51.54 | 6.77 [0.08 – 13.46] | 10, 5.76 | -15.26 [-23.13 – -7.39] | 42, 3.29 | -0.13 [-0.27 – 0.02] |
| Epetraborole-omadacycline-ethambutol | 3.02 [2.36 - 3.68] | 7, 57 | 7.85 [5.54 – 10.16] | 10, 7.53 | -16.89 [-16.94 - -16.83] | 42, 11.5 | 0.10 [-0.07 – 2.27] |
| Ceftriaxone-omadacycline-rifabutin | 3.02 [1.68 - 4.35] | 7, 57 | 7.87 [1.52 - 14.23] | 10, 6.34 | -9.88 [-40.16 - 20.43] | 42, 5.64 | -0.07 [-0.25 – 0.11] |
| Azithromycin-rifabutin-ethambutol | 3.02 [0 - 6.83] | 7, 11.06 | 1.04 [-0.84 – 2.92] | 10, 6.34 | -2.45 [-4.28 - -0.62] | 42, 2.62 | -0.04 [-0.12 – 0.05] |
| Non-treated controls | 3.02 [2.12 - 3.41] | 42, 1.29 | -0.05 [-0.07 - -0.02] | - | - | - |  |

*Italics* means slope 95% confidence crossed zero, and thus a flat horizontal line with slope equal to zero. Brackets are 95% confidence intervals.

### Microbial kill and rebound based on CFU/mL readout

For the CFU/mL readout, microbial kill results during the 42 days of treatment were as shown in **Figure 3B**. **T**he largest amplitude for microbial kill below *B_0_* for each regimen was compared to that of the SOC, on the anti-log scale. The mean (± standard deviation) fold-change kill below *B_0_*compared to SOC were 595±362.2-fold for azithromycin-minocycline-ethambutol, 281.6±256.8-fold for the azithromycin-omadacycline-ethambutol regimen, 206.7±10.09-fold for ceftriaxone-omadacycline-rifabutin, and 604.0±58.91-fold for the epetraborole-omadacycline-ethambutol regimen, but these values did not statistically differ from each other (Kruskal Wallis *p*=0.32).

In **Figure 3B**, the bacterial burden pattern changes were described by two BFs, shown in **Table 4**. BF_1_ was best described by the ODE 𝜸 kill slope. For context, non-treated control had no BF_1_ (𝜸=0 throughout the study), while the SOC BF_1_ was described by 𝜸=0.16 log_10_ CFU/mL/day until hinge 1 at 7 days. **Table 4** shows that fastest kill speed was ceftriaxone-omadacycline-rifabutin, with a 𝜸=0.55 log_10_ CFU/mL/day with end of microbial kill [hinge 1] at 21 days and residual bacterial burden of 1.59 log_10_ CFU/mL on that day. This means that the ceftriaxone-omadacycline-rifabutin regimen killed MAC 3.44-times faster than the SOC and killed bacteria in the HFS-MAC with a greater amplitude that the SOC. **Figure 3B** shows BF_2_s for the rebound growth. The *𝜀* for each regimen is shown in **Table 4**. For context, non-treated controls had a BF_2_ *𝜀* of 0.09 log_10_ CFU/mL/day for the wild-type total MAC burden versus 0.15 log_10_ CFU/mL/day for the SOC. The *𝜀* for all combinations were statistically similar to each other, and to the non-treated controls, which meant the resistant population grew as fast as wild type.

**Table 4.**
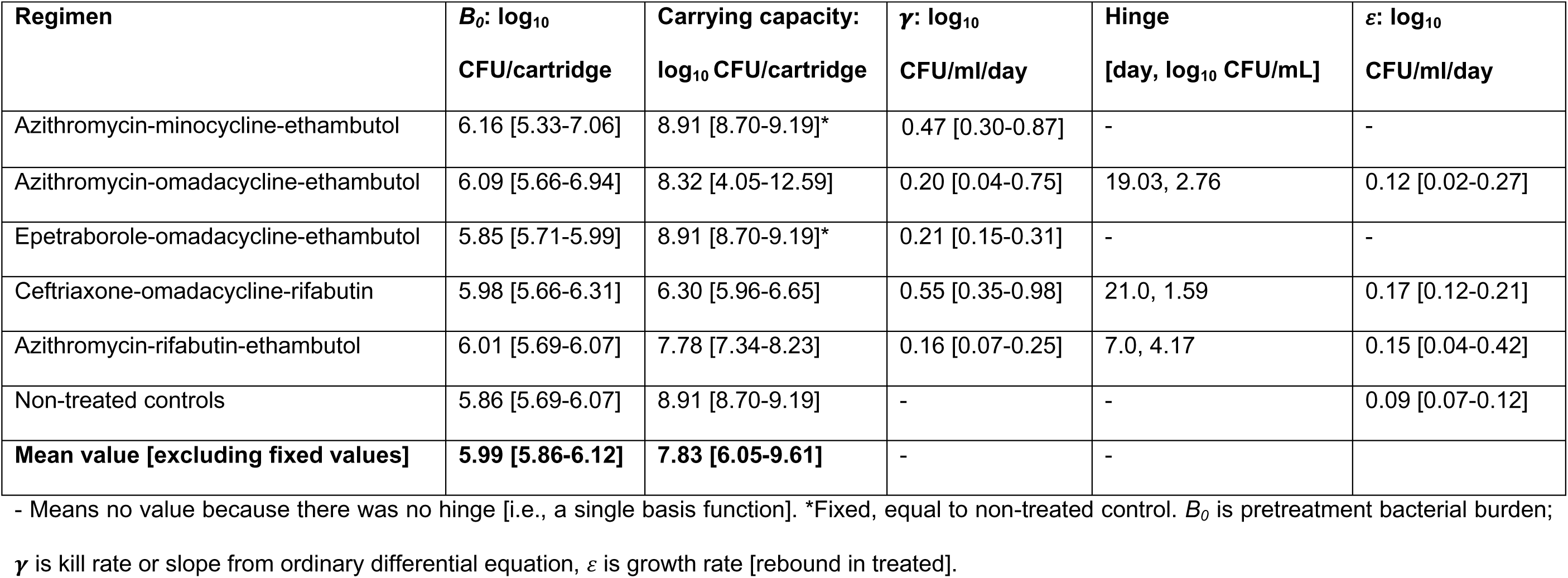
Parameter Estimates for microbial dynamics (CFU) on agar and 95% Credible Intervals for BF_1_ and BF_2_.

| Regimen | $B_0$: $\log_{10}$<br>CFU/cartridge | Carrying capacity:<br>$\log_{10}$ CFU/cartridge | $\gamma$: $\log_{10}$<br>CFU/ml/day | Hinge<br>[day, $\log_{10}$ CFU/mL] | $\varepsilon$: $\log_{10}$<br>CFU/ml/day |
| --- | --- | --- | --- | --- | --- |
| Azithromycin-minocycline-ethambutol | 6.16 [5.33-7.06] | 8.91 [8.70-9.19]* | 0.47 [0.30-0.87] | - | - |
| Azithromycin-omadacycline-ethambutol | 6.09 [5.66-6.94] | 8.32 [4.05-12.59] | 0.20 [0.04-0.75] | 19.03, 2.76 | 0.12 [0.02-0.27] |
| Epetraborole-omadacycline-ethambutol | 5.85 [5.71-5.99] | 8.91 [8.70-9.19]* | 0.21 [0.15-0.31] | - | - |
| Ceftriaxone-omadacycline-rifabutin | 5.98 [5.66-6.31] | 6.30 [5.96-6.65] | 0.55 [0.35-0.98] | 21.0, 1.59 | 0.17 [0.12-0.21] |
| Azithromycin-rifabutin-ethambutol | 6.01 [5.69-6.07] | 7.78 [7.34-8.23] | 0.16 [0.07-0.25] | 7.0, 4.17 | 0.15 [0.04-0.42] |
| Non-treated controls | 5.86 [5.69-6.07] | 8.91 [8.70-9.19] | - | - | 0.09 [0.07-0.12] |
| **Mean value [excluding fixed values]** | **5.99 [5.86-6.12]** | **7.83 [6.05-9.61]** | - | - |  |

- Means no value because there was no hinge [i.e., a single basis function]. \*Fixed, equal to non-treated control. $B_0$ is pretreatment bacterial burden; $\gamma$ is kill rate or slope from ordinary differential equation, $\varepsilon$ is growth rate [rebound in treated].

### Inflection in microbial kill curves versus resistance emergence in BF_2_

Changes in the drug-resistant subpopulations (azithromycin, ethambutol, rifabutin, epetraborole, minocycline) by regimen were as shown **Supplementary Figure S3** for the drug-resistant burden expressed as log_10_ CFU/mL. **Figure 4** shows the same results but based on resistant subpopulation as % of the total population as CFU/mL. Further results are discussed in **Supplementary data** and shown in **Supplementary Figure S3A-F.**

**Figure 4.**
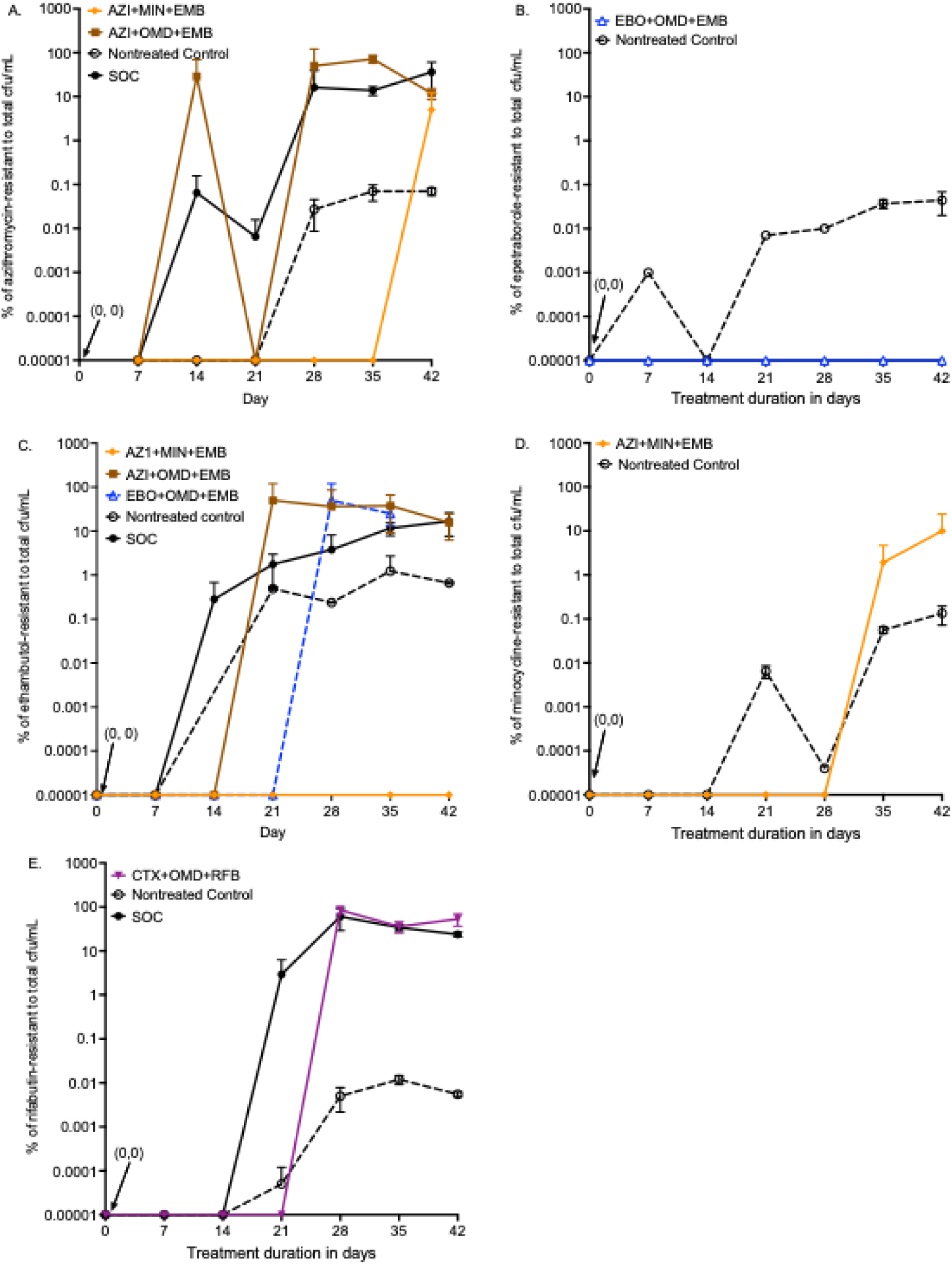
Emergence of drug resistance within each combination regimen as % of total bacterial burden. Symbols are means and error bars are standard deviation. **A.** AZI=Azithromycin. **B.** EBO=Epetraborole. **C.** EMB=Ethambutol. **D.** MINO=Minocycline. **E.** RFB=Rifabutin

**Figure 4** shows that for all drugs, % drug-resistant subpopulation in the non-treated remained below 0.1%. Any drug-resistant subpopulation % greater than un non-treated controls was considered resistance amplification. Regarding azithromycin-resistance, **Figure 4A** shows rapid amplification by both the SOC and azithromycin-omadacycline-ethambutol regimens, while azithromycin-minocycline-ethambutol delayed that amplification. **Figure 4B** shows that the epetraborole-omadacycline-ethambutol regimen completely suppressed epetraborole-resistance. In **Figure 4C**, the azithromycin-minocycline-ethambutol suppressed emergence of ethambutol-resistance while other regimens amplified ethambutol-resistance. In **Figure 4D**, azithromycin-minocycline-ethambutol amplified minocycline-resistance. In **Figure 4E**, ceftriaxone-omadacycline-rifabutin amplified rifabutin-resistance. Regarding omadacycline and ceftriaxone, which degrade in agar in the face of slow growth of MAC, we did not capture the drug-resistant subpopulation size.

The drug-resistant subpopulations were modeled using the logistic function, with results shown in **Supplementary Table S1**. We compared the logistic function model parameters of each drug-resistant subpopulation to that of the total rebound bacterial burden, with results shown in **Table 5**. For the SOC, the azithromycin-resistant population logistic growth curve versus that of the total population had an r^2^=0.66 (p=0.2), that for rifabutin-resistant was an r^2^=0.27 (p=0.32), while that for the ethambutol-resistant subpopulation had an r^2^>0.99 (p=0.01). In essence BF_2_ for the SOC rebound was statistically identical to that of the ethambutol-resistant subpopulation. This means that ethambutol-resistance was the most explanatory of rebound growth, and not azithromycin (r^2^<0.7). For the ceftriaxone-omadacycline-rifabutin regimen, while the p=0.08 did not reach statistical significance, the r^2^=0.7 for rifabutin versus rebound was considered high and needs further investigation.

**Table 5.** Rebound growth rate of regimen [BF2] versus drug-resistant subpopulation growth.

| **Regimen and drug-resistance** | **r<sup>2</sup>** | **p-value** |
| --- | --- | --- |
| **Standard of care** | Referent | Referent |
| Azithromycin-resistant | 0.66 | 0.20 |
| Rifabutin-resistant | 0.27 | 0.32 |
| Ethambutol-resistant | >0.99 | **0.01** |
| **Azithromycin-omadacycline-ethambutol** | Referent | Referent |
| Azithromycin-resistant | 0.44 | 0.17 |
| Ethambutol-resistant | 0.49 | 0.15 |
| **Ceftriaxone-omadacycline-rifabutin** | Referent | Referent |
| Rifabutin-resistant | 0.7 | 0.08 |
| **Epetraborole-omadacycline-ethambutol** |  |  |
| Epetraborole | - | - |
| Ethambutol | - | - |

-No resistance or no BF<sub>2</sub>

## DISCUSSION

Here, we introduced the concept of BFs for modeling bacterial exposure-response surfaces as the pattern changes from microbial kill to rebound (12). BFs were originally introduced by Paul de Casteljau and Pierre Bézier working on the problems of car design (12, 21–23). Here, we adopted BFs for MAC microbial kill and resistance emergence. First, framing the relationship as BFs allows testing the hypothesis of how much a particular drug-resistant subpopulation is explanatory of the total rebound MAC population. This gives a quantitative tool to test for the effect of resistance to one drug versus failure of a combination. Second, the BFs differed by readout. It was assumed that liquid culture fluorescent detection of oxygen during mycobacterial metabolism in the automated MGIT liquid assay is as accurate, if not more sensitive and more precise at lower bacterial burdens, than CFU counts of agar (24, 25). Moreover, given the different dynamic range in MGIT time-to-positivity (57 days) versus CFU/mL on agar (0 to approximately 1 billion CFU/mL), this means that growth curves derived using solid agar cannot be imposed on those from liquid cultures (unless first translated), and the BFs for microbial kill and growth, must be independently derived for solid and liquid cultures. In fact, there are multiple assays that measure different dimensions of bacterial death, with different dynamic ranges (26). BFs give an approach to integrate data from all these readouts, without arbitrarily assuming superiority of one over another. Third, elsewhere, we have demonstrated that multivariate adaptive regression splines (an artificial intelligence algorithm) output of BFs and equations automatically enables multiscale systems modeling for multiple readouts, starting from the level of cellular chemical reactions, to organs, to whole body physiological parameters and maturation, and populations of patients (12, 13, 27). This means that the bacterial response BFs we described here from two readouts can be integrated to human virtual models of disease progression, to be considered with BFs for drug metabolism, to model cure an toxicity rates of drugs (27).

The main purpose of the study was systematic PK/PD-based screening to find a combination regimen with faster sterilizing effect than the SOC. To this end, based on both the TTP and CFU/mL readouts, the fastest kill speed was by ceftriaxone-omadacycline-rifabutin, at 3.44-7.57 times the SOC. The extent of kill below *B_0_*, and kill speed of this regimen will need to be confirmed in up to 5 MAC isolates to model heterogeneity, based on our recommendations elsewhere (6). Moreover, the ceftriaxone dose in a multidrug combination needs to be optimized. Here, we used a ceftriaxone peak of 44.65 mg/L which is below the peak serum concentrations for 2G a day of 280 mg/L, and given the 12.18-fold penetration into ELF below a peak of 3,380 mg/L in the lung, that is only 1.3% of intrapulmonary concentrations (28, 29). Moreover, the finding that rifabutin-resistance curves were explanatory (r^2^=0.7) of the regimen’s rebound growth means that rifabutin is likely why ceftriaxone-omadacycline-rifabutin had rebound growth. Given that tedizolid was the second highest ranked drug in the quantitative analysis, it will be important to test if replacing rifabutin with tedizolid abolishes resistance amplification (6). Another possible strategy is to administer ceftriaxone-omadacycline-rifabutin till the end of the kill slope (hinge), after which it is followed by the runner up regimen of azithromycin-minocycline-ethambutol to complete therapy. These strategies can be investigated in the future HFS-MAC studies.

We determined the role of SOC drug components in failure of the regimen, using the BF approach. Meta-analyses of prospective clinical studies have shown that macrolide-containing regimens achieved better sustained sputum culture conversion rates than non-macrolide regimens as initial therapy (6). Retrospective clinical studies have demonstrated that macrolide-based regimens without ethambutol or rifamycin had higher risk of development of macrolide-resistance (3–5). However, in previous HFS-MAC study of dual azithromycin and rifabutin regimen, failure of therapy arose in parallel with the emergence of acquired ethambutol-resistance (30). Here, we found that the same was true even in the triple combination (azithromycin-ethambutol-rifabutin) – there was emergence of resistance to all three drugs, but the one that drove the shape of the rebound (BF_2_) and was most explanatory of the rebound, was ethambutol and not azithromycin. On the other hand, we have also shown that ethambutol on its own does not kill below *B_0_;* and while macrolides killed below *B_0_* but the amplitude was poor (9). This suggests that likely ethambutol is the Achilles heel of SOC because development of ethambutol-resistance leads to failure of the entire SOC. Moreover, since ceftriaxone-omadacycline-rifabutin outperformed azithromycin containing regimens, there is likely nothing intrinsically superior about macrolides and likely achieved their superior reputation because of drugs they were historically compared to.

Finally, even though this was MAC study, the methods we used here should be straightforward to adapt to anti-infectives for non-mycobacterial infectious. First, comparisons of antibiotics by speed and amplitude compared to the SOC for different infections could add to the current approaches that rely on such metrics as 1.0 log and 2.0 log_10_ CFU/mL kill. It answers the different question – will a regimen kill faster and more extensively than the current SOC, and suppress resistance emergence better than the SOC. Second, the ODE obviates the need to force microbial kill patterns into prespecified “straightjacket” patterns such as exponential decline which were historically derived from a limited number of readouts (26). Third, the BFs can be used to model microbial kill rates and resistance emergence to any antimicrobial, as well as to compare the speeds of kill and rebound, rapidly growing bacteria, fungi, viruses, and even protozoa. Since the BFs are building blocks for constructing more complex functions, they allow building a single complex function (*f*_C_) for any antimicrobial or combination thereof, for the entire duration of an experiment.

Our study has some limitations, as captured by QC in **Table 1**. First, we did not use multiple MAC isolates, since this was just a screening study. Second, we did not employ factorial design (9). As a result, the ceftriaxone dose in the combination needs further optimization. Third, we did not translate the 𝜸 slope to sustained sputum conversion in patients’ sputa. Fourth, Monte Carlo experiments were not performed in this screening study. These limitations will be addressed in studies that have already commenced.

In summary, we screened azithromycin- and non-azithromycin combinations for drugs chosen based on being able to kill MAC below *B_0_*. We used the mathematics of basis functions for analysis. Ceftriaxone-omadacycline-rifabutin achieved the fastest speed of kill, and highest kill below *B_0_*. The regimen will need further optimization of the ceftriaxone dose, and possible replacement of rifabutin with tedizolid.

## METHODS

### Bacterial isolates, materials, and MICs

The standard laboratory strain of *Mycobacterium avium* (ATCC#700898) and THP-1 monocyte cell line (ATCC#TIB-202) were purchased from the American Type Culture Collection. Phorbol myristate acetate (PMA), Roswell Park Memorial Institute (RPMI) 1640 medium, and heat-inactivated fetal bovine serum were purchased from Sigma-Aldrich (St. Louis, MO). Hollow-fiber cartridges (catalog number C7011; polyvinylidene difluoride) were purchased from FiberCell (Frederick, MD, USA). MAC cultures were propagated in Middlebrook 7H9 broth supplemented with 10% oleic acid, albumin, dextrose, and catalase (OADC), whereas CFU/mL was determined using Middlebrook 7H10 agar supplemented with 10% OADC was propagated from stock cultures in all the experiments. MGIT system and supplies were purchased from Becton-Dickinson (USA). TTP data was collected from the MGIT system using the Epicenter Software. MICs of each drug were determined using the standard broth microdilution method (31).

### Comparing different combination regimens in the HFS-MAC

The construction of the HFS-MAC has been extensively described in the past (16, 18–20, 32–47). Construction includes modeling *B_0_* of MAC lung disease in patients which is a median is 5.17 [range: 4.23-6.2] log_10_ CFU/mL in cavities and 3.0 (range: 0.48-3.85) log_10_ CFU/mL in nodular/bronchiectatic lesions, intracellular infection in monocyte-lineage cells, and intrapulmonary PKs of drugs (48–50). Each HFS-MAC unit was inoculated with 20 mL of ATCC#700898 MAC infected THP-1 monocytes, and drug treatment started after 24h of inoculation of the HFS-MAC units.

Drugs were administered once a day via computer-controlled syringe pumps, for 42 days (6 weeks). The following combinations were tested in the HFS-MAC: Regimen 1 (R1) comprising azithromycin plus minocycline plus ethambutol, R2 comprising of azithromycin plus ethambutol plus omadacycline, R3 comprising of epetraborole plus ethambutol plus omadacycline, and R4 of ceftriaxone plus omadacycline plus rifabutin, R5 was the SOC comprised of azithromycin plus rifabutin plus ethambutol, and R6 was non-treated controls. The intra-pulmonary PK properties intended for the HFS-MAC study are shown in **Supplementary Table S2**, together with MICs. The lung PK parameters were based on previously published work (28, 29, 51–62). There were two HFS-MAC units per regimen. The central compartment of each HFS-MAC unit was sampled at steady state on day 7, at 0 (pre-dose), 1, 4, 8, 12, 16, 23.5h post-dosing. The peripheral compartment of each HFS-MAC unit was sampled for bacterial burden estimation on day 0, 3, 7, 10, 14, 21, 28, 35, and 42. Samples were inoculated on plain 7H10 agar, as well as agar supplemented with three-time the MIC concentration of azithromycin or epetraborole or ethambutol or minocycline or rifabutin. Omadacycline and ceftriaxone resistance determination was not performed due to the rapid drug degradation at the incubation temperature. Total bacterial burden was also assessed by inoculating samples into the MGIT assay tubes to record the TTP (days).

### Quality Score

We have introduced a quality score for pre-clinical MAC PK/PD HFS-MAC models (9). The tool has 6 scoring criteria. The score is used to assess the quality and adequacy of the experiment, and for adequacy of PK/PD design (9). We applied this to our work here. Other QC data were %CV of CFU/mL, TTP, and drug concentration measurements on sampling days

## ACKNOWLEDGEMENT

Shashikant Srivastava thank Dr. Jullie V Philley for discussions that led to design of this study and partial funding support to perfume the initial studies.

## ETHICAL APPROVAL

Not applicable.

## AUTHOR CONTRIBUTIONS

Conceptualization and design: Shashikant Srivastava (SKS) and PJM. MIC and HFS-MAC experiments: SS, GDB, and SKS. PK/PD modeling, basis functions, and application of ODEs: TG. TG wrote the first draft of the manuscript. All authors reviewed and approved the final version of the manuscript for publication.

